# Use of a Sibling Subtraction Method (SSM) for Identifying Causal Mutations in *C. elegans* by Whole-Genome Sequencing

**DOI:** 10.1101/166355

**Authors:** Braveen B. Joseph, Nicolas A. Blouin, David S. Fay

**Author notes:** Department of Molecular Biology, College of Agriculture and Natural Resources, University of Wyoming, Laramie, WY 82071. University of Wyoming INBRE Bioinformatics Core (Nicolas A. Blouin).

## Abstract

Whole-genome sequencing (WGS) has become an indispensible tool for identifying causal mutations obtained from genetic screens. To reduce the number of causal mutation candidates typically uncovered by WGS, *C. elegans* researchers have developed several strategies. One involves crossing N2-background mutants to the polymorphic strain CB4856, which can be used to simultaneously identify mutant-strain variants and obtain high-density mapping information. This approach, however, is not well suited for uncovering mutations in complex genetic backgrounds, and CB4856 polymorphisms can alter phenotypes. Several other approaches make use of DNA variants introduced by mutagenesis. This information is used to implicate genomic regions with high densities of DNA lesions that persist after mutant backcrossing, but these methods provide lower mapping resolution than use of CB4856. To identify suppressor mutations using WGS, we developed a new approach termed the Sibling Subtraction Method (SSM). This method works by eliminating variants present in both mutants and their non-mutant siblings, thus greatly reducing the number of candidates. We used this method here with two members of the *C. elegans* NimA-related kinase family, *nekl-2* and *nekl-3*. Combining weak aphenotypic alleles of *nekl-2* and *nekl-3* leads to penetrant molting defects and larval arrest. We isolated ~50 suppressors of *nekl-2; nekl-3* synthetic lethality using F1-clonal screening methods and a toxin-based counter-selection strategy. When applied to five of the identified suppressors, SSM led to only one to four suppressor candidates per strain. Thus SSM is a powerful approach for identifying causal mutations in any genetic background and provides an alternative to current methods.

## Introduction

Whole-genome sequencing (WGS) was first used to identify causal mutations in *C. elegans* nearly 10 years ago (SARIN *et al*. 2008), and this approach has been progressively refined and applied to a growing number of organisms (SCHNEEBERGER AND WEIGEL 2011; LESHCHINER *et al*. 2012; OBHOLZER *et al*. 2012; JAMES *et al*. 2013; MORESCO *et al*. 2013; HU 2014; SCHNEEBERGER 2014; DOITSIDOU *et al*. 2016). One complication of WGS is that many hundreds or thousands of variants are typically detected in individual mutant strains, thus making it difficult to pinpoint the causal mutation that is responsible for the observed phenotype. Because the large majority of mutations that alter phenotypes lead to changes in the primary sequence of proteins (SARIN *et al*. 2008), filtering steps can be applied such that only DNA variants that alter coding sequences are considered. Nevertheless, in the absence of other information, this can still result in large numbers of exonic variants that must be experimentally tested.

Several WGS strategies have been developed to reduce the number of candidate variants by simultaneously providing mapping data on the causal mutation, an approach termed “mapping by sequencing” (SCHNEEBERGER AND WEIGEL 2011; SCHNEEBERGER 2014; DOITSIDOU *et al*. 2016). In *C. elegans*, a widely used approach makes use of the Hawaiian (HA) variant, CB4856, which differs from the field-standard N2 (Bristol) strain by >100,000 polymorphisms (CONSORTIUM 1998; WICKS *et al*. 2001; HILLIER *et al*. 2008; DOITSIDOU *et al*. 2010; VERGARA *et al*. 2014). Typically, mutants generated in the N2 background are crossed to the HA strain to generate N2/HA hybrids, which are then allowed to self-fertilize. Phenotypically mutant progeny of N2/HA hermaphrodites are then isolated and subjected to WGS and variant identification. N2/HA polymorphisms can then be exploited to map mutations to genomic regions that are homozygous for N2-specific variants, thereby greatly reducing the number of causal variants to be considered (DOITSIDOU *et al*. 2010). Variations on HA mapping have also been developed to facilitate the mapping and identification of diverse allele types (SMITH *et al*. 2016). This approach, however, has two major caveats. One is that it is not easily applied to complex genotypes, such as mutations that alter phenotypes only when one or more additional mutant loci are present in the background. In addition, genetic differences between N2 and HA can lead to changes in the expression of phenotypes in ways that are not predictable (DOROSZUK *et al*. 2009; REDDY *et al*. 2009; BENDESKY *et al*. 2012; RODRIGUEZ *et al*. 2012; GREEN *et al*. 2013; POLLARD AND ROCKMAN 2013; GLATER *et al*. 2014; SNOEK *et al*. 2014; DOITSIDOU *et al*. 2016; KAMKINA *et al*. 2016; SINGH *et al*. 2016).

As an alternative, the method known as EMS Density Mapping does not require use of the HA strain and relies on the ability to detect signature changes to DNA that are induced by the commonly used mutagen ethyl methanesulfonate (EMS) (DRAKE AND BALTZ 1976; FLIBOTTE *et al*. 2010; SARIN *et al*. 2010). After serial backcrossing, strains are subjected to WGS, and genomic regions that contain higher densities of EMS lesions, in particular those flanking causal mutations, are identified (ZURYN *et al*. 2010). Because the density of EMS-induced lesions is relatively low compared with the number of polymorphisms between N2 and HA, this method yields substantially lower mapping resolution than the HA method (DOITSIDOU *et al*. 2016).

Another approach that circumvents use of the HA strain, the Variant Discovery Method, also relies on variant frequencies to identify chromosomal regions of interest but uses both EMS and non-EMS variants (MINEVICH *et al*. 2012; CHEESMAN *et al*. 2016). Thus, the Variant Discovery Method may improve mapping resolution relative to that of EMS Density Mapping but provides less information than HA mapping methods (DOITSIDOU *et al*. 2016).

We describe here the Sibling Subtraction Method (SSM), which is an alternative approach for WGS analysis that does not require use of the HA strain and does not conceptually rely on variant mapping methods. We applied this strategy to identify suppressors of synthetically lethal *nekl-2; nekl-3* double mutants, which arrest as larvae because of defects in molting (YOCHEM *et al*. 2015; LAZETIC AND FAY 2017). Our results indicate that this method can be used to reduce the number of candidate causal variants to as few as one or two in most cases, thus providing a powerful alternative to current approaches. This study highlights the utility of using synthetically lethal combinations of weak aphenotypic alleles as a genetic background for suppressor screening and includes a description of a counter-selection approach to increase the efficiency of genetic suppressor screens.

## Materials and Methods

### Strains and maintenance

*C. elegans* strains were maintained according to standard protocols (STIERNAGLE 2006) and were propagated at 22°C unless otherwise stated. Strains used in this study include N2/Bristol (wild type), WY1145 [*nekl-2(fd81); nekl-3(gk894345); fdEx286* (pDF153, *nekl-3(+);* pTG96, SUR-5::GFP)] (LAZETIC AND FAY 2017), WY1208 [*nekl-2(fd81); nekl-3(gk894345); fd130*], WY1209 [*nekl-2(fd81); fd131; nekl-3(gk894345)*], WY1210 [*nekl-2(fd81); fd132; nekl-3(gk894345)*], WY1211 [*nekl-2(fd81); fd133; nekl-3(gk894345)*], WY1217 [*nekl-2(fd81); nekl-3(gk894345); fd139*], and WY1255 [*nekl-2(fd81); nekl-3; fdEx303* (pDF153, *nekl-3(+);* pTG96, SUR-5::GFP, PMA122, P_*hsp16.41:peel-1*_)] (SEIDEL *et al*. 2011). For additional strains, see Table S1.

### Variant detection

Paired-end libraries were prepared for all strains and sequenced on an Illumina HiSeq 2000. Resulting reads were analyzed *via* SSM as described in the Results. A detailed description of sample preparation and variant analysis is provided in the Supplemental WGS and SSM Methods section. Briefly, our workflow was adapted from CloudMap and incorporated tools including Trimmomatic, BWA, SAMtools, SnpEff, SnpSift, Genome Analysis Toolkit, and MPileup Integrative Genomics Viewer VarScan, implemented on the UseGalaxy.org platform (LI AND DURBIN 2009; LI *et al*. 2009; MCKENNA *et al*. 2010; CINGOLANI *et al*. 2012a; CINGOLANI *et al*. 2012b; KOBOLDT *et al*. 2012; MINEVICH *et al*. 2012; BOLGER *et al*. 2014; AFGAN *et al*. 2016).

### RNAi

dsRNA injections and RNAi feeding were carried out according to standard protocols (AHRINGER 2005). dsRNA was prepped using PCR products with T7 polymerase sequences; for details on primers used for these studies, see the Supplemental Genetics Methods section.

### Transgene rescue

Rescue experiments were carried out according to standard protocols (EVANS AND HUNTER 2005). For details on fosmids and plasmids used, see the Supplemental Genetics Methods section.

*CRISPR*. CRISPR/Cas9 editing was carried out on strain WY1145 using *dpy-10* Co-CRISPR methods in combination with RNP injection protocols (ARRIBERE *et al*. 2014; PAIX *et al*. 2015). For details on guide and repair template RNAs used, see the Supplemental Genetics Methods section.

## Results and Discussion

To better understand the functions of Nek family kinases during development, we sought to identify extragenic suppressors of molting defects in *nekl-2* and *nekl-3* mutants. Previous studies had identified an allelic series of *nekl-2* and *nekl-3* mutations including null alleles that arrest at the L1/L2 molt, moderate loss-of-function alleles that arrest as L2/L3s, and weak alleles that are aphenotypic (YOCHEM *et al*. 2015; LAZETIC AND FAY 2017). Furthermore, certain combinations of weak alleles of *nekl-2* and *nekl-3* lead to double mutants that display penetrant molting defects and concomitant larval arrest. In the case of strain WY1145, synthetically lethal *nekl-2(fd81); nekl-3(gk894345)* double mutants are maintained by the presence of a rescuing extrachromosomal array (*fdEx286*) that is transmitted to ~65% of self-progeny and expresses wild-type *nekl-3* along with a fluorescent reporter (SUR-5::GFP) (LAZETIC AND FAY 2017) (and data not shown). Thus, in this study, WY1145 hermaphrodites gave rise to viable array-containing GFP^+^ progeny as well as array-negative GFP^−^ progeny, 98.8% (n = 444) of which were arrested as larvae with molting defects (Figure 1A). Rare GFP^−^ escapers that reached adulthood produced progeny that arrested nearly uniformly with molting defects (98.5%, n=1721).

**Figure 1.**
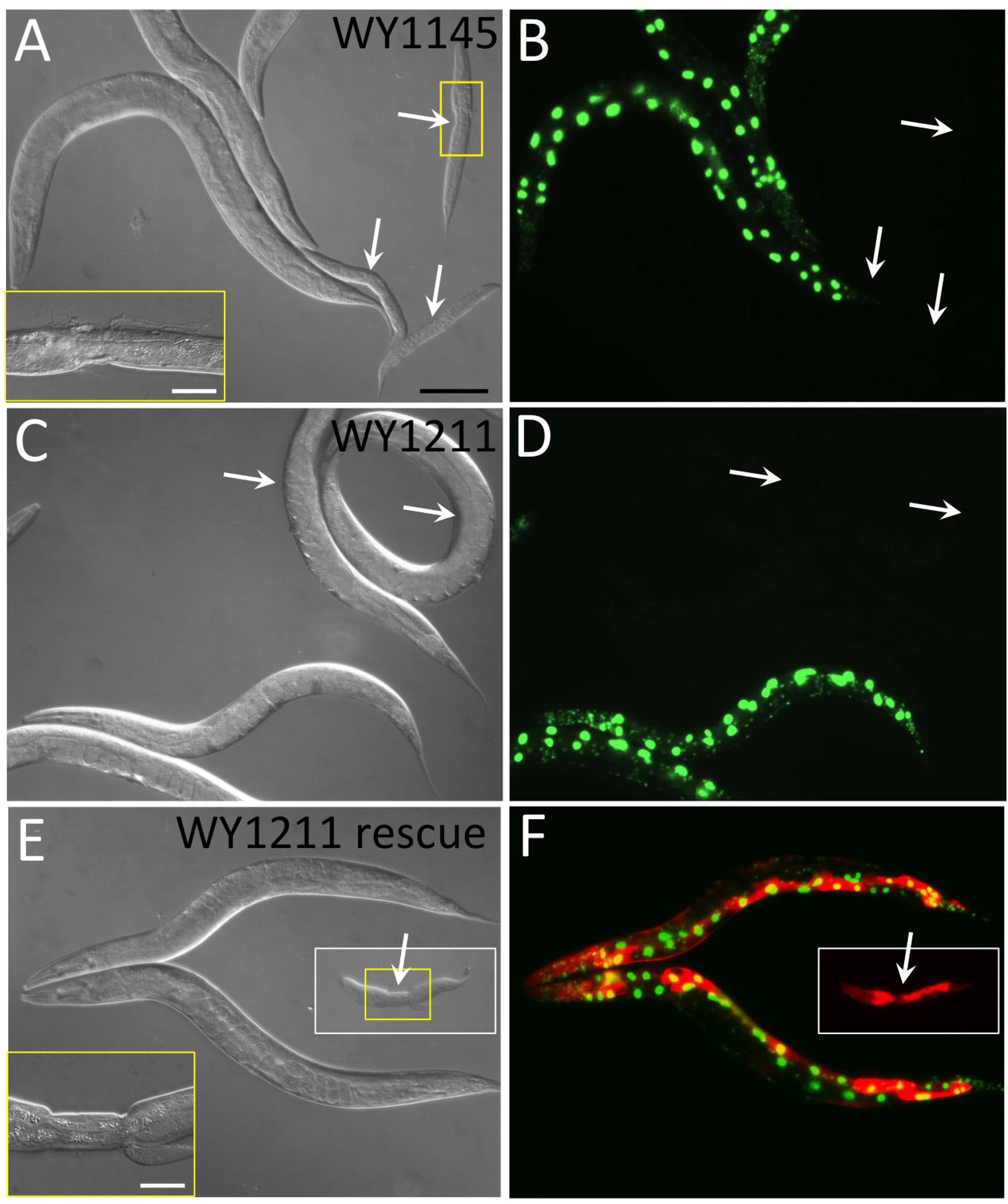
Phenotypes of molting-defective and suppressed strains. (A,C,E) DIC and (B,D,F) fluorescence images of strains WY1145 [*nekl-2(fd81); nekl-3(gk894345)*] and WY1211 [*nekl-2(fd81); paa-1(fd134); nekl-3(gk894345)*]. GFP^+^ worms carry an extrachromosomal array (*fdEx286*) that expresses wild-type *nekl-3* and SUR-5::GFP. (A,B) Whereas GFP^−^ worms (white arrows) in the starting strain, WY1145, arrest uniformly with molting defects, (C,D) GFP^−^ worms in the suppressed strain, WY1211, can reach adulthood. (E,F) Suppression is reverted in strain WY1211 by the expression of wild-type *paa-1* from a fosmid, which is carried by an independent extrachromosomal array marked with SUR-5::RFP. Compare GFP^−^ RFP^+^ arrested larva (E,F) with GFP^−^ gravid adults (C,D). Also note that the *paa-1^+^* RFP-marked array is not generically deleterious for growth as evidenced by the viability of GFP^+^ RFP^+^ adults (E,F; also see Supplementary Genetics Methods). White arrows indicate GFP^−^ worms; white boxes in (E,F) indicate that the image was acquired from a region outside the main panel; yellow boxes indicate regions of increased magnification. Black scale bar in A (for A–F) = 100 µm; white scale bars in insets (A,E) = 20 µm.

We reasoned that WY1145 could be effective for identifying extragenic suppressors because (1) both *fd81* and *gk894345* are very weak loss-of-function alleles and thus are potentially amenable to genetic suppression, and (2) by carrying out the screen with double mutants, we could obtain suppressors that are specific to either *nekl-2* or *nekl-3*, as well as suppressors of both. As a first approach, we mutagenized WY1145 with EMS and carried out a semi-clonal F1 screen (Figure S1). Plates containing candidate suppressor mutations were initially identified based on the increased frequency of F2 GFP^−^ animals reaching adulthood and were subsequently verified by propagating strains derived from isolated F2 GFP^−^ animals for multiple generations. From this screen of ~8000 haploid genomes, 27 independent isolates were obtained with suppression frequencies ranging from ~20% to nearly 90% (Figure 1 and Table S1). Notably, a pilot screen using a moderate loss-of-function allele, *nekl-3(sv3)*, alone failed to uncover strong suppressors (data not shown), suggesting that approaches using synthetic-lethal combinations of weak alleles may be particularly productive.

To reduce the labor associated with F1 clonal/semi-clonal screening methods, we considered alternative strategies. Because the frequency of naturally occurring GFP^−^ escapers reaching adulthood in WY1145 (~1–2%) was likely to be much higher than the frequency of authentic suppressors, standard non-clonal approaches were deemed impractical because of the high incidence of false positives. We therefore considered a strategy involving a counter-selectable marker to eliminate animals carrying the *nekl-3^+^* rescuing array. *peel-1* is a sperm-derived toxin that when overexpressed in *C. elegans* larvae and adults leads to death at all postembryonic stages (SEIDEL *et al*. 2008; SEIDEL *et al*. 2011). We engineered a *nekl-2(fd81); nekl-3(gk894345)* double-mutant strain (WY1255) with a rescuing array containing wild-type *nekl-3*, SUR-5::GFP, and a heat shock–inducible *peel-1* transgene. After an incubation of WY1255 at 34°C for 2 hr, nearly 100% of array-containing GFP^+^ worms perished within 24 hr. As outlined in Figure S2, WY1255 worms were mutagenized, and 100 individual P0 animals were picked to large plates and incubated at 22°C until F2 progeny began to emerge (~5–6 days). Plates were then heat shocked to kill array-containing animals and were monitored for an additional 7–8 days to identify plates with actively propagating populations of GFP^−^ animals. To prevent the re-emergence of any surviving GFP^+^ worms, plates were heat shocked again after 4 days. From this screen we identified an additional 23 suppressors, demonstrating the utility of this counter-selection approach (Table S1). In particular, this method should be useful for identifying suppressors of phenotypes that are <100% penetrant, but it can also be applied to phenotypes that are fully penetrant.

Suppressed strains (*nekl-2; nekl-3; sup*) were backcrossed to WY1145 and, after preliminary genetic characterization, five strains were selected for further analysis by WGS (see Supplemental Genetics Methods for backcrossing and other genetics methods). These mutants encompassed a range of suppressor strengths as well as dominant and recessive alleles (Table S1; Figure 1C,D). In addition, genetic analyses determined that three of these five suppressors were on LGX, whereas two were autosomal (Table S1). The molecular identification of the suppressors, however, presented several technical challenges given that the suppressor mutations did not exhibit obvious phenotypes on their own. In addition, the *nekl-2(fd81)* and *nekl-3(gk894345)* mutations are aphenotypic and show defects only when combined as double mutants (LAZETIC AND FAY 2017). Although it was possible to introduce both the *fd81* and *gk894345* mutations into the HA mapping strain (CB4846) using CRISPR/Cas9 methods, as discussed in the Introduction, this divergent background has the potential to alter the expression or penetrance of phenotypes. Furthermore, other methods that do not require the use of the HA strain may not provide sufficient mapping resolution of the affected loci to consistently enable easy identification of causal mutations (DOITSIDOU *et al*. 2016).

We therefore devised a WGS strategy to identify *nekl-2; nekl*-*3* suppressors that did not depend on either variant density mapping or the use of CB4856. We refer to our approach as the Sibling Subtraction Method (SSM), and a generic version of this strategy is shown in Figure 2; a detailed description of our approach is provided in the Supplemental WGS and SSM Methods section. Importantly, this strategy can be applied to a variety of phenotypes as well as both simple and complex genetic backgrounds, including mutations that lead to enhancement or suppression. To test this approach, *nekl-2(fd81); nekl-3(gk894345)* suppressed strains (*nekl-2; nekl-3; sup*) were crossed to WY1145 to generate heterozygous (*sup/+*) GFP^+^ F1 cross-progeny, which were picked to individual plates and allowed to self-fertilize (Figure 2A and Figure S3). Next, sibling GFP^+^ F_2_ progeny from a single F_1_ parent were picked to individual plates and allowed to produce the F_3_ generation. Based on standard Mendelian inheritance patterns, a 1:2:1 ratio corresponding to *sup/sup, sup/+*, and *+/+* would be expected among the F_2_ generation. Plates containing F_2_ parents of the *sup/sup* genotype were recognized based on the high frequency of viable GFP^−^ adult F_3_s. Moreover, viable GFP^−^ F_3_ worms from candidate *sup/sup* plates were picked to new plates and allowed to propagate to ensure suppressor homozygosity. Conversely, plates containing parental F_2_s of the *+/+* genotype were identified by the absence of suppressed (adult GFP^−^) F_3_ animals (Figure S3). For each suppressor strain, individual *sup/sup* isolates were combined to make genomic DNA, which we refer to as the “Suppressed” DNA pool (Figure S3). Corresponding *+/+* isolates were also combined to generate a “Non-Suppressed Sibling Comparator” DNA pool (Figure S3). We note that the number of independent *sup/sup* or *+/+* isolates used to generate the DNA pools ranged from 5–20 (average = 11), indicating that as few as 5 isolates may be sufficient for our approach to be successful.

**Figure 2.**
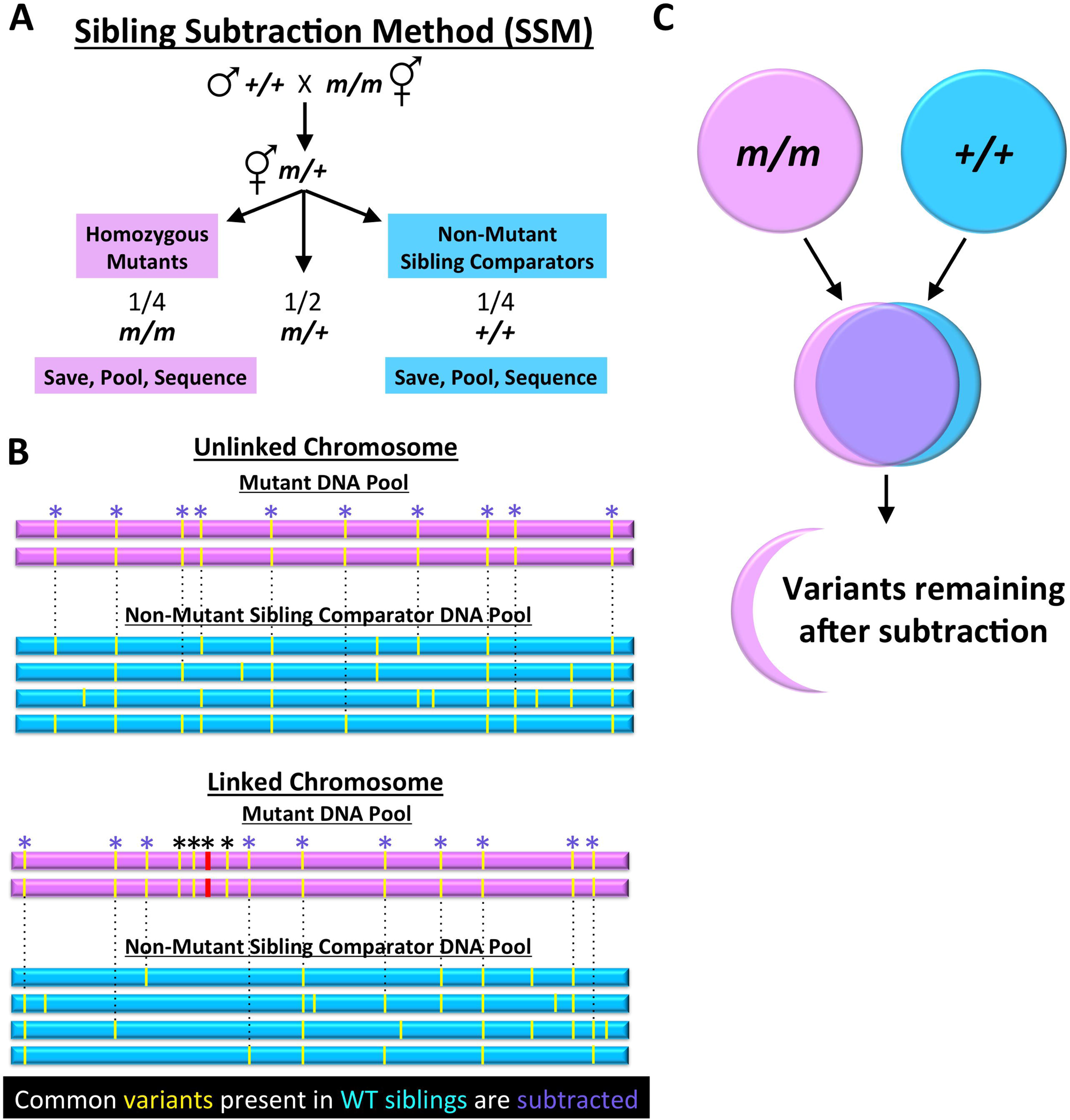
Schematic design of the Sibling Subtraction Method. (A) The genetic crosses required for generating multiple independent *m/m* and sibling *+/+* isolates, which are combined to produce the Mutant DNA pool and the Non-Mutant Sibling Comparator DNA pool, respectively (also see Figure S3 and Supplemental Genetics Methods). (B) Simplified representations of sequenced chromosomes from the Mutant pool (pink) and Non-Mutant Sibling Comparator pool (blue) are shown. Yellow lines indicate variants detected by WGS, and asterisks indicate variants that are homozygous in the Mutant pool. Note that the depiction underestimates the true number of variants per chromosome. Dashed lines and purple asterisks indicate variants that are present in both the Mutant Pool and the Non-mutant Sibling Comparator Pool, which can be eliminated (subtracted) as candidate causal mutations. Note that subtraction will remove most or all variants on unlinked chromosomes, whereas homozygous variants very close to the causal mutation (red) may not be subtracted (black asterisks). (C) Venn diagram of subtracted variants (purple) along with the relatively small proportion of remaining candidate variants (pink) after application of the SSM.

A workflow for the experimental and bioinformatical analysis is shown in Figure 3 and is described in detail in the Supplemental Methods sections. In our analysis of WGS/SSM data, two criteria were initially applied for variants to be considered candidate causal mutations. First, variants must be homozygous (present in 100% of reads) in the *sup/sup* DNA pool. Second, variants must be completely absent (present in 0% of reads) in the corresponding *+/+* sibling DNA pool. By using these two criteria, we predicted that the large majority of strain-specific non-causal mutations acquired during mutagenesis and/or subsequent propagation would be subtracted. In addition, any variants present in the starting strain that differed from the published wild-type sequence would also be subtracted (Figure 2B,C). In practice, we found that this subtraction step eliminated 93.4% (range, 91–98%) of the homozygous variants detected in the five Suppressed DNA pools (Table 1). Likewise, when considering homozygous variants present in the suppressed strains that led to predicted changes in the resulting protein (nonsense, missense, and splice site mutations), we found that an average of 97.1% (range, 96– 98%) of variants were subtracted. Finally, after manual filtering of the predicted variant reads, we obtained an average of 2.2 candidate causal variants per strain (range, 1–4), with four of five strains having two or fewer candidates (Table 1; Supplemental File 1).

**Figure 3.**
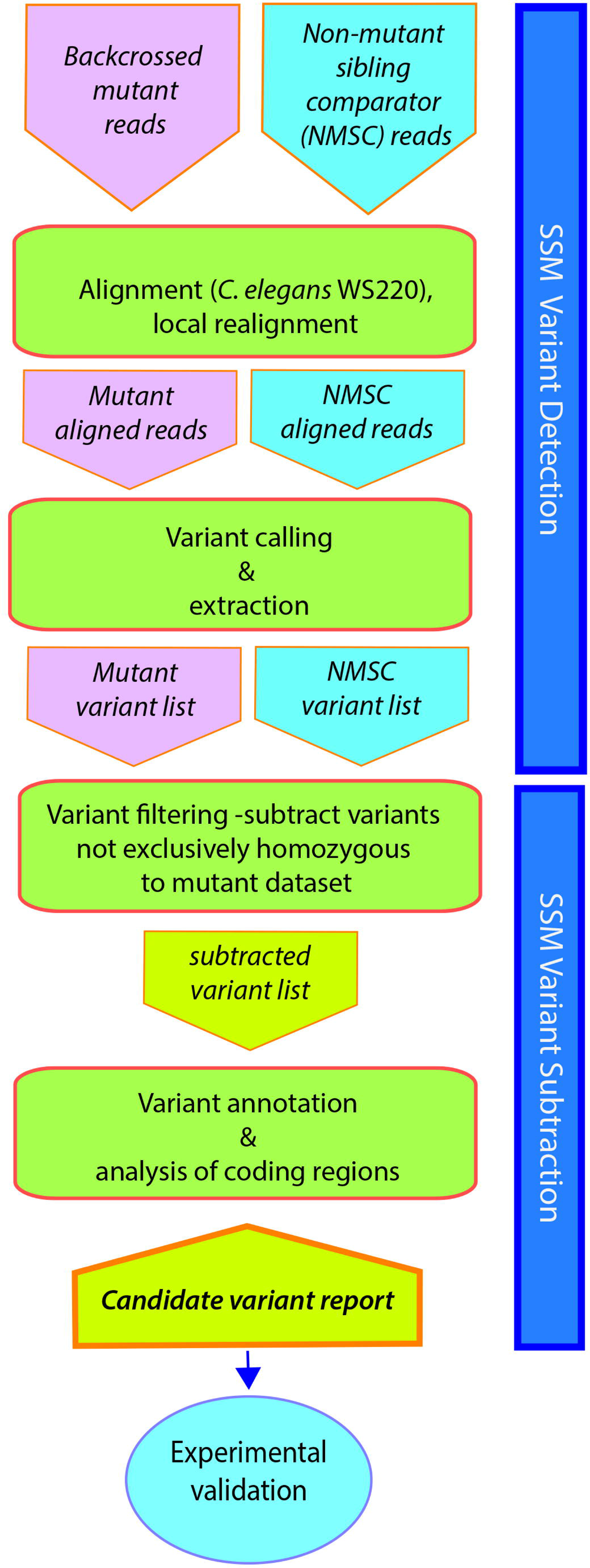
SSM workflow. Overview of the experimental and bioinformatical steps involved in the SSM/WGS method. For additional details, see text and Supplemental Methods sections.

We first validated the identities of all four recessive causal mutations using a combination of RNAi, transgenic rescue experiments, and CRISPR/Cas9 (see Supplemental Genetics Methods). One example is strain WY1211 (*nekl-2; nekl-3; fd134*), which exhibits 34% suppression (n = 199), making it the weakest of the suppressors subjected to SSM/WGS (Figure 1). After subtraction, only two candidate variants remained that affected coding regions, corresponding to missense mutations in C27F2.9 and F48E8.5/*paa-1* (Table 1). Notably, both variants reside within a 0.6-map-unit-long region on the left arm of LGIII. C27F2.9 encodes a protein with homology to JMJD8, and the detected variant leads to a R220C substitution in a residue that is not highly conserved among closely related species. *paa-1* encodes an essential gene that is orthologous to the PR65 structural subunit of the mammalian PP2A protein phosphatase. Moreover, the identified variant in *paa-1* causes a G550E substitution in a glycine residue that is highly conserved throughout metazoans (data not shown). Two lines of evidence demonstrated that the *fd134* causal mutation corresponds to the lesion in *paa-1*. First, we recreated the identical (G→A) mutation using CRISPR/Cas9 methods in the WY1145 background and obtained two independent suppressed lines. Second, we injected fosmids encoding the wild-type *paa-1* locus into WY1211 and observed clear de-suppression in three of three lines (Figure 1[**E**,**F]**). Our identification of *paa-1* as a suppressor of *nekl-2; nekl-3* defects suggests that the *C. elegans* PP2A complex may oppose NEKL-2/3 kinase activity, possibly by dephosphorylating NEKL2/3 targets.

In the case of the dominant suppressor (*fd132*; strain WY1210), WGS/SSM identified coding variants in two genes, T09B9.4 (M399I) and W07E11.1 (P1181L). Notably, both genes are located on LGX, which genetic mapping had shown to harbor the causal mutation, and are positioned within ~300,000 bp of each other (~0.25 map units). Moreover, both genes reside within a 1,704,753-bp region that contains 13 consecutive variants that are present in 100% of Suppressed DNA reads and 0% of the Non-Suppressed Sibling Comparator reads. Genomic engineering of the T09B9.4 and W07E11.1 mutations using CRISPR/Cas9, however, did not lead to genetic suppression, indicating that neither mutation likely corresponds to the causative change in this strain (also see Supplemental Genetics Methods). Thus, the *fd132* causal mutation may reside in a non-coding region, such as an element that promotes transcriptional or post-transcriptional repression, or WGS/SSM may have failed to identify the causal coding variant (also see below).

We next compared our subtraction approach to the EMS density mapping method using the data generated from our Suppressed sequencing reads (ZURYN *et al*. 2010). As shown in Figure S4, although a subset of peaks was consistent with the locations of our identified causal mutations, EMS density mapping methods did not in most cases provide unambiguous resolution of the causal mutant loci. In contrast, the SSM strategy was successful in limiting the number of high-probability (genetic code–altering) candidates to as few as one or two loci (Table 1).

**Table 1.** Summary of data from SSM.

| Strain | Variants before Subtraction |  |  |  | Variants after Subtraction |  |  |
| --- | --- | --- | --- | --- | --- | --- | --- |
|  | Suppressed |  | Non-Suppressed<br>Sibling Comparators |  | Total<sup>a</sup> | Δ Coding<sup>a</sup> | Δ Coding<br>Filtered<sup>b</sup> |
|  | Total<sup>a</sup> | Δ Coding<sup>a</sup> | Total<sup>a</sup> | Δ Coding<sup>a</sup> |  |  |  |
| WY1208 | 2317 | 214 | 2324 | 216 | 112 | 7 | 4 |
| WY1209 | 2316 | 224 | 2391 | 220 | 173 | 9 | 2 |
| WY1210 | 2563 | 222 | 2315 | 222 | 212 | 4 | 2 |
| WY1211 | 2330 | 233 | 2364 | 217 | 59 | 4 | 2 |
| WY1217 | 2534 | 225 | 2339 | 215 | 243 | 8 | 1 |

<sup>a</sup> Numbers correspond to data derived prior to manual filtering of sequence reads. Also see Supplemental WGS and SSM Methods section and Supplemental File 1.
<sup>b</sup> Numbers obtained after manual filtering of reads.

We also varied our subtraction criteria to determine how this would affect the number of candidate variants. In our initial analysis we required candidate variants to be present in 100% of reads from the Suppressed DNA pool and to be completely absent from the Non-Suppressed Sibling Comparator pool. However, experimental error or sequencing artifacts could potentially eliminate authentic causal mutations, giving rise to false negatives. For example, DNA pools prepared from dominant alleles (e.g., *fd132*) could contain animals that were heterozygous, thus reducing the percentage of causal variant reads in the Suppressed DNA pool to <100%. Conversely, in the case of weakly penetrant alleles, heterozygous strains could be accidentally assigned to the Non-Suppressed Sibling Comparator pool, thus increasing the percentage of causal variant reads to >0%.

We therefore reanalyzed data from strains WY1209 and WY1210 using three different filtering parameters. First, we reduced the requirement for Suppressed DNA reads from 100% to ≥90%, while maintaining Non-Suppressed Sibling Comparator reads at 0% (Table S2 and Supplemental File 1). For strain WY1209, this increased the total number of variants after subtraction by only 3% and did not increase the number of variants affecting coding regions. For strain WY1210, the number of variants after subtraction increased by only 6% but included three changes affecting coding regions: R03E5.10 (M536I), T05A10.1 (P1628S), and F53A9.7 (G29E) contained 34/36 (94%), 14/15 (93%), and 2/23 (91%) mutant variant reads in the Suppressed DNA pool, respectively. Thus, the *fd132* causal mutation may affect a non-coding region or one of the three additional genes implicated by the relaxed filtering criteria.

Reducing the cutoff for the Non-Suppressed Sibling Comparator pool to ≤10% of total variant reads while maintaining the stringency of the Suppressed DNA pool at 100% led to 1% and 4% increases in the number of variants after subtraction for strains WY1209 and WY1210, respectively (Table S2 and Supplemental File 1). Finally, relaxing both criteria (≥90% of Suppressed pool reads and ≤10% of Comparator pool reads) resulted in 5% and 12% increases in the number of variants after subtraction for strains WY1209 and WY1210, respectively (Table S2 and Supplemental File 1). This led to a total of three variants that affected coding regions in WY1209 and five variants in WY1210, versus two for each strain using the original (100%:0%) filtering criteria. Thus, relaxing both filtering parameters by 10% did not strongly alter the efficiency of variant subtraction by SSM and may reduce the occurrence of false negatives due to experimental error or sequencing artifacts.

In summary, we find SSM/WGS to be a simple and potent approach for identifying causal mutations in *C. elegans* and should be applicable to other model systems. Importantly, this approach does not require the use of polymorphic strains (e.g., CB4856) and does not rely on variant mapping methods. We note that although SSM/WGS appears to provide better resolution than EMS density mapping based on our analysis (Figure S4), our approach did not allow for a direct comparison between SSM and “bulked segregant” Variant Discovery Method methods; future parallel assessments of these methods may prove useful. In addition, we showed that a counter-selection method using an extrachromosomal array encoding an inducible toxin can provide a highly efficient means for identifying genetic suppressors, especially in cases of incomplete penetrance of the starting strain. In addition, our studies underscore the utility of using weak alleles in synthetic-lethal combinations as starting strains for suppressor screens. In particular, recent advances in genomic engineering (ARRIBERE *et al*. 2014; PAIX *et al*. 2014; PAIX *et al*. 2015; DICKINSON AND GOLDSTEIN 2016), as well as the Million Mutation resource (THOMPSON *et al*. 2013), make the identification of weak alleles and combinatorial-dependent phenotypes highly feasible. Finally, because SSM/WGS is not impacted by genetic background, it should be possible to identify genetic modifiers of complex genotypes in a straightforward manner.

## Acknowledgements

Some strains were provided by the CGC, which is funded by NIH Office of Research Infrastructure Programs (P40 OD010440). This work was supported by NIH grants GM066868 (to DSF), P20 GM103432 (Wyoming INBRE), and P20 GM103451 (New Mexico INBRE, bioinformatics core). We also thank Amy Fluet for editing.

