## Supplementary Materials for "Use of a Sibling Subtraction Method (SSM) for Identifying Causal Mutations in *C. elegans* by Whole-Genome Sequencing"

**Table S1. List of *nekl-2; nekl-3* suppressors.**

| Strains isolated from F1 semi-clonal screen | | | | |
|---|---|---|---|---|
| Strain Name | Allele | % Suppression (n) | Dominant/Recessive | Autosome/LGX |
| Strains subjected to SSM & WGS | | | | |
| WY1208 | *fd130* | 83 (223) | Recessive | LGX |
| WY1209 | *fd131* | 87 (360) | Recessive | Autosome |
| WY1210 | *fd132* | 50 (276) | Dominant | LGX |
| WY1211 | *fd133* | 35 (422) | Recessive | Autosome |
| WY1217 | *fd139* | 76 (235) | Recessive | LGX |
| Strains NOT subjected to SSM & WGS | | | | |
| WY1212 | *fd134* | 34 (387) | Weak Dominant | N.D. |
| WY1213 | *fd135* | N.D. | N.D. | N.D. |
| WY1214 | *fd136* | 25 (374) | Weak Dominant | N.D. |
| WY1215 | *fd137* | 35 (277) | Weak Dominant | N.D. |
| WY1216 | *fd138* | 48 (315) | N.D. | N.D. |
| WY1218 | *fd140* | 47 (430) | N.D. | N.D. |
| WY1220 | *fd142* | 37 (138) | N.D. | N.D. |
| WY1221 | *fd143* | 33 (425) | N.D. | N.D. |
| WY1222 | *fd144* | 33 (167) | N.D. | N.D. |
| WY1223 | *fd145* | 32 (613) | N.D. | N.D. |
| WY1224 | *fd146* | 31 (30) | N.D. | N.D. |
| WY1225 | *fd147* | 30 (389) | N.D. | N.D. |
| WY1267 | *fd151* | 63 (300) | Recessive | Autosome |
| WY1268 | *fd152* | 55 (77) | N.D. | N.D. |
| WY1269 | *fd153* | 79 (333) | N.D. | N.D. |
| WY1270 | *fd154* | 51 (206) | Recessive | Autosome |
| WY1271 | *fd155* | 94 (306) | Recessive | LGX |
| WY1272 | *fd156* | N.D. | N.D. | N.D. |
| WY1273 | *fd157* | N.D. | N.D. | N.D. |
| WY1274 | *fd158* | N.D. | N.D. | N.D. |
| WY1275 | *fd159* | N.D. | N.D. | N.D. |
| WY1276 | *fd160* | N.D. | N.D. | N.D. |
| Strains isolated from non-clonal F1-clonal counter-selection screen | | | | |
| Strains NOT subjected to SSM & WGS | | | | |
| WY1277 | *fd161* | 63 (260) | N.D. | N.D. |
| WY1278 | *fd162* | 78 (639) | Recessive | Autosome |
| WY1279 | *fd163* | 83 (121) | N.D. | N.D. |
| WY1280 | *fd164* | N.D. | N.D. | N.D. |
| WY1281 | *fd165* | 57 (130) | N.D. | N.D. |
| WY1282 | *fd166* | 34 (938) | Recessive | Autosome |
| WY1283 | *fd167* | 71 (219) | N.D. | N.D. |
| WY1284 | *fd168* | N.D. | N.D. | N.D. |

| | | | | |
|---|---|---|---|---|
| WY1285 | *fd169* | 86 (179) | N.D. | N.D. |
| WY1286 | *fd170* | 43 (112) | Recessive | Autosome |
| WY1287 | *fd171* | 77 (220) | N.D. | N.D. |
| WY1288 | *fd172* | 74 (282) | N.D. | N.D. |
| WY1289 | *fd173* | 13 (643) | Recessive | Autosome |
| WY1290 | *fd174* | 77 (137) | N.D. | N.D. |
| WY1291 | *fd175* | 85 (66) | N.D. | N.D. |
| WY1292 | *fd176* | 78 (201) | N.D. | N.D. |
| WY1293 | *fd177* | N.D. | N.D. | N.D. |
| WY1294 | *fd178* | 46 (190) | N.D. | N.D. |
| WY1295 | *fd179* | 84 (266) | N.D. | N.D. |
| WY1296 | *fd180* | 76 (223) | N.D. | N.D. |
| WY1297 | *fd181* | 80 (93) | N.D. | N.D. |
| WY1298 | *fd182* | 30 (123) | Recessive | Autosome |
| WY1299 | *fd183* | N.D. | N.D. | N.D. |
