## Supplementary Materials for "Use of a Sibling Subtraction Method (SSM) for Identifying Causal Mutations in *C. elegans* by Whole-Genome Sequencing"

**Table S2. Comparison of the standard and alternative workflows and filtering criteria.**

| Subtraction Criteria | WY1209 Variants[a] | WY1210 Variants[a] |
|---|---|---|
| **Standard workflow[b]** | | |
| 100% Mutant:0% NMSC | 173 (2) | 212 (2) |
| **Alternative workflow[b]** | | |
| 100% Mutant:0% NMSC | 193 (2) | 146 (2) |
| ≥90% Mutant:0% NMSC | 199 (2) | 154 (5) |
| 100% Mutant:≤10% NMSC | 194 (4) | 151 (2) |
| ≥90% Mutant:≤10% NMSC | 201 (4) | 163 (5) |

[a] Numbers in parentheses indicate manually filtered variants that affect coding regions. Also see Supplemental File 1.

[b] Standard and alternative workflows are described in the Supplemental WGS and SSM Methods section. Because the specific computational tools used for these two analyses necessarily differed, the number of subtracted variants for 100% Mutant:0% NMSC (Non-Mutant Sibling Comparator) also differs between the two workflows.
