## Supplementary Materials for "Use of a Sibling Subtraction Method (SSM) for Identifying Causal Mutations in *C. elegans* by Whole-Genome Sequencing"

### Supplemental Genetics Methods

***Backcrossing and initial assessment of genetic traits***. We strongly recommend backcrossing of all strains obtained from genetic screens. Moreover, phenotypic assessment should not be carried out on strains that have not been backcrossed ~5× to avoid drawing incorrect conclusions caused by background mutations. In the case of mapping and WGS, straightforward backcrossing procedures can be used to determine if the mutation is stable, behaves as a single Mendelian locus, is on LGX or an autosome, and is dominant or recessive.

Hermaphrodites from suppressed strains were backcrossed to WY1145 [*nekl-2* (*fd81*); *nekl-3* (*gk894345*); *fdEx286* (*nekl-3*$^{+}$ + SUR-5::GFP)] males as shown below.

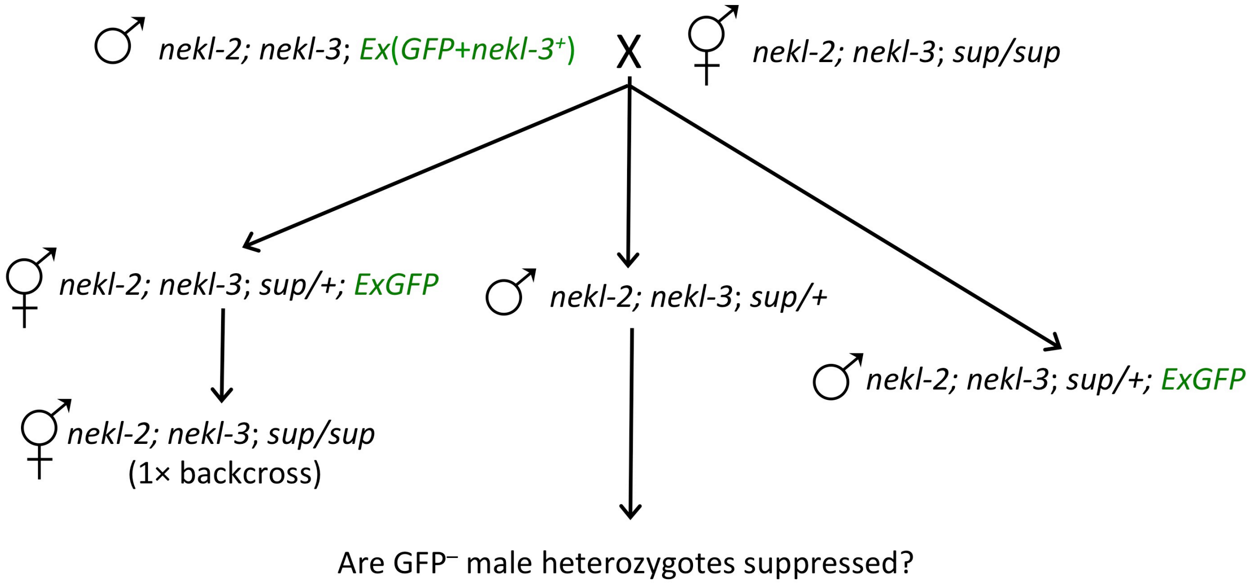


GFP$^{+}$ F1s were picked to individual plates and allowed to self-fertilize to produce suppressed (*sup/sup*) F2s. In addition, adult F1 males were scored to determine whether they contained the rescuing array—GFP$^{+}$ versus GFP$^{-}$. The presence of GFP$^{-}$ males indicates that the suppressors are either dominant, on LGX, or both. Representative results for these backcrosses are shown below.

| Strain | GFP$^{-}$ ♂ | GFP$^{+}$ ♂ | n = | Frequency GFP$^{-}$ ♂ |
|---|---|---|---|---|
| WY1145 | 0 | 92 | 92 | 0.00 |
| WY1208 | 50 | 46 | 96 | 0.52 |
| WY1209 | 0 | 65 | 65 | 0.00 |
| WY1210 | 26 | 56 | 82 | 0.31 |
| WY1211 | 1 | 70 | 71 | 0.01 |
| WY1212 | 8 | 32 | 40 | 0.20 |
| WY1214 | 5 | 48 | 53 | 0.09 |
| WY1215 | 18 | 43 | 61 | 0.13 |
| WY1217 | 7 | 55 | 62 | 0.11 |

Frequency = ♂ / (♂ + ♂)

The parental strain, WY1145, serves as a control. The above data indicate that the suppressors in WY1209 (*fd131*) and WY1211 (*fd133*) are fully recessive and autosomal, whereas the suppressors in WY1208 (*fd130*) and WY1210 (*fd132*) may be dominant or on LGX; other suppressors, such as WY1217 (*fd139*) may also be dominant/semi-dominant or on LGX. In addition to these tests, ~50 GFP$^{+}$ F2s were picked for each cross and allowed to self. If suppression is due to a single altered locus, we would expect to observe relatively strong suppression (*sup/sup*) from 25% of the F2 plates, based on the presence of GFP$^{-}$ F3s. For all the strains we tested (WY1208–WY1212 and WY1217), we observed relatively high levels of suppression at frequencies consistent with a single suppressor locus (range, 14–38%; n = 50 for each strain). A similar frequency of F2 plates failed to show any suppression, as expected, for the +/+ isolates (also see Figure 2 and Figure S3).

***Analysis of dominance versus recessiveness.*** To further analyze the properties of the suppressors, we generated strain WY1232 [*nekl-2* (*fd81*); *nekl-3* (*gk894345*); *fdEx186* (*nekl-3*$^{+}$ + SUR-5::GFP); *fdEx197* (SUR-5::RFP)], which carries a second extrachromosomal array marked by RFP. This strain allows for the unambiguous identification of cross-progeny hermaphrodites as shown below.

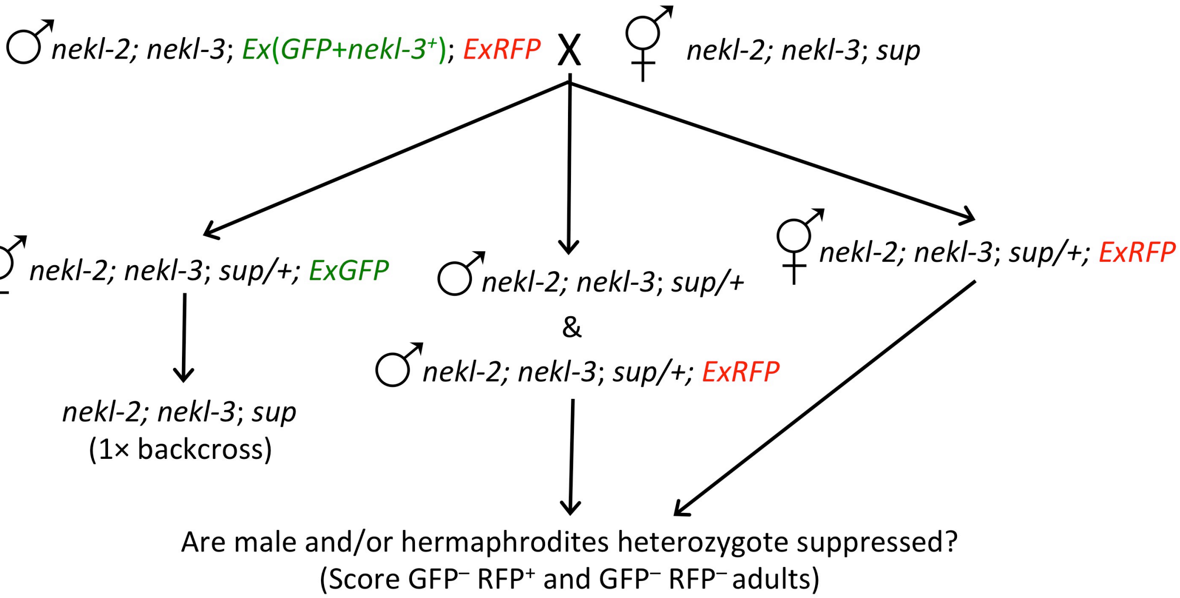


We tested F1 cross-progeny hermaphrodites for suppression based on the presence of RFP$^{+}$ GFP$^{-}$ males and hermaphrodites, as well as RFP$^{-}$ GFP$^{-}$ males. Representative results for males are shown below, which are consistent with findings from crosses to WY1145 males.

| Strain | 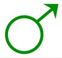 | 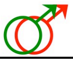 | 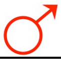 | 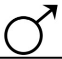 | n = | Freq. 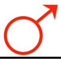 |
|---|---|---|---|---|---|---|
| WY1208 | 18 | 25 | 50 | 9 | 102 | 0.67 |
| WY1209 | 31 | 35 | 0 | 1 | 67 | 0.00 |
| WY1210 | 11 | 26 | 20 | 4 | 61 | 0.44 |
| WY1211 | 21 | 16 | 1 | 0 | 38 | 0.06 |
| WY1212 | 7 | 8 | 5 | 0 | 23 | 0.38 |
| WY1214 | 28 | 16 | 3 | 0 | 47 | 0.16 |
| WY1215 | 10 | 26 | 15 | 2 | 54 | 0.37 |
| WY1217 | 22 | 33 | 7 | 0 | 62 | 0.18 |

Frequency = 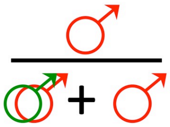

Representative results for hermaphrodites are shown below. These indicate that the WY1208 suppressor is recessive, and thus on LGX, whereas the WY1210 suppressor is dominant. These results also suggest that the WY1217 suppressor may be recessive and on LGX.

| Strain | 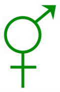 | 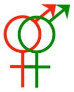 | 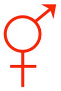 | n = | Freq. |
|---|---|---|---|---|---|
| WY1208 | 54 | 8 | 0 | 62 | 0.0 |
| WY1209 | 34 | 20 | 1 | 55 | 0.05 |
| WY1210 | 50 | 16 | 11 | 77 | 0.41 |
| WY1211 | 43 | 11 | 1 | 55 | 0.08 |
| WY1214 | 46 | 27 | 6 | 79 | 0.19 |
| WY1215 | 26 | 7 | 4 | 37 | 0.36 |
| WY1217 | 36 | 12 | 1 | 49 | 0.08 |

Frequency = 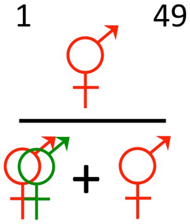

To determine if the dominant suppressor mutation in WY1210 is on LGX or is autosomal, we carried out crosses as outlined below.

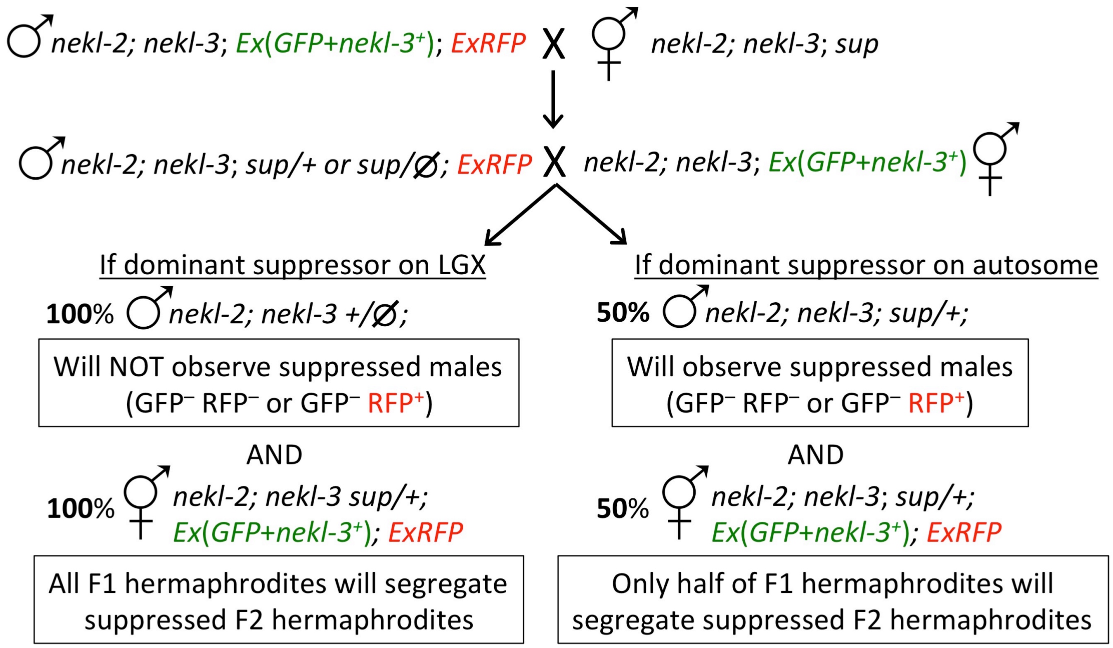


Whereas we observed >100 GFP$^+$ RFP$^-$ and/or GFP$^+$ RFP$^+$ males resulting from the second cross, we failed to observe any GFP$^-$ RFP$^+$ or GFP$^-$RFP$^-$ males, indicating linkage to LGX. Furthermore, 100% (n = 40) of GFP$^+$RFP$^+$ hermaphrodites from the second cross produced suppressed progeny, confirming the location of *fd132* on LGX.

***Confirmation of causal suppressor mutations.***

*Transgene rescue experiments*. Standard transgenic rescue using a wild-type copy of the causal mutant gene typically leads to correction of the mutant phenotype in worms that carry the transgene. In the case of suppressor mutations, rescue leads to de-suppression and therefore expression of the mutant phenotype (e.g., molting defects). Two approaches were used to test for rescue of candidate suppressor variants. In one approach, wild-type copies of candidate genes present on fosmids were injected with SUR-5::GFP into *nekl-2; nekl-3; sup* adults, and GFP$^+$ F1 progeny were scored for molting defects. In some cases, incomplete rescue (de-suppression) led to our ability to score progeny from the F2 generation, although we were unable to obtain stably transmitting lines. For strain WY1208, we injected fosmids corresponding to candidate C04A11.4 (WRM0620dD12, WRM0632aG02, and WRM0610cA04, 2–6 ng/µl each + SUR-5::GFP [pTG96], 100 ng/µl). Suppression by *fd130* in WY1208 led to 83% viability (17% arrest; Table S1). After injection, 80% of GFP$^+$ F1 larvae (n = 25) arrested with molting defects, and a similar trend was observed in the F2 progeny from several viable transmitting F1s (83% arrest, n = 37). In contrast, injection of this fosmid mix into wild type showed no deleterious effects on larval development or adult viability. Likewise we injected WY1217 with fosmids corresponding to candidate B0302.1 (WRM0610cD03 and WRM0612dE01, 6 ng/µl each + SUR-5::GFP [pTG96], 100 ng/µl). Whereas the *fd139* mutation led to 76% viability in this strain (24% arrest), we observed arrest in 93% of GFP$^+$ F1s (n = 30).

For the weaker suppressor strain WY1211 (*fd133*), we took an alternative approach to more firmly establish rescue. We injected a mix of fosmids corresponding to F48E8.5 and an RFP marker (WRM0618bG08, WRM0640bA06, WRM0618aG08, ~6 ng/µl each + SUR-5::RFP [pTG96.2], 100 ng/µl ) into WY1211 worms that also carried the *fdEx286 nekl-3*$^{+}$ GFP$^{+}$ rescuing array (Figure 1C,D). In this way, we were able to establish three stably transmitting GFP$^{+}$ RFP$^{+}$ lines and then scored viability in the F3s. Arrest among the GFP$^{-}$ RFP$^{+}$ animals occurred at frequencies of 92% (n = 210), 95% (n = 128), and 97% (n = 181) for the three lines tested, respectively, whereas arrest among GFP$^{-}$ RFP$^{-}$ worms was only 63% (n = 223; p < 0.0001) (Figure 1E,F). Furthermore, <10% of GFP$^{+}$ RFP$^{+}$ progeny underwent arrest (n > 200 for each strain), indicating that the arrays were not highly toxic (Figure 1E,F).

*RNAi.* dsRNAs corresponding to exons in candidate suppressor genes were prepared using standard PCR methods followed by T7 RNA synthesis. These were injected at 0.8–1.0 µg/µl into strain WY1145 (Ahringer 2005). GFP$^{-}$ F1 progeny were scored, together with non-injected controls, for each experiment. In the case of WY1217, injection of dsRNA corresponding to exon 11 of B0302.1 (1033-bp fragment) led to 18.6% (n = 1112) adult viability versus 0.7% (n = 279) in non-injected controls (p < 0.0001). For WY1209, injection of dsRNA corresponding to exons 9–11 of F56D12.6a (970-bp fragment) led to 16.5% (n = 1054) adult viability versus 1.9% (n = 210) in non-injected controls (p < 0.0001).

*CRISPR/Cas9.* CRISPR/Cas9 methods were used to generate genetic lesions in top suppressor candidates for strains WY1208 (C04A11.4), WY1209 (F56D12.6), WY1210 (T09B9.4 and W07E11.1), WY1211 (F48E8.5), and WY1217 (B0302.1). Specifically, CRISPR/Cas9 RNPs were injected into strain WY1145 using *dpy-10* co-CRISPR methods, and Rol and Dpy progeny were monitored for suppression (Arribere *et al.* 2014; Paix *et al.* 2014; Paix *et al.* 2015). In cases where mutations in the candidate suppressors led to the premature termination of transcripts (C04A11.4, F56D12.6, and B0302.1), we used CRISPR/Cas9 to generate breaks within 45 bp of the detected lesion and allowed non-homologous repair mechanisms to generate new stop codons or frameshifts. This led to the isolation of three new alleles of C04A11.4 (*fd208*, 1-bp deletion; *fd209*, 2-bp deletion; *fd210*, 8-bp deletion), two new alleles of F56D12.6 (*fd211*, 13-bp deletion; *fd212*, 10-bp deletion), and three new alleles of B0302.1 (*fd213*, 16-bp insertion; *fd214*, 7-bp deletion; *fd215*, 4-bp insertion). All identified CRISPR suppressor alleles led to premature translational termination and strong suppression of WY1145 larval arrest. Together with RNAi and rescue data, these results confirmed the identities of the causal mutations in WY1208, WY1209, and WY1217 strains.

For the essential gene F48E8.5, we used a repair template to introduce the lesion identified by WGS and also introduced a site for the DraI (TTTAAA) endonuclease. This led to the isolation of two independent alleles (*fd216* and *fd217*), both of which contained the identical (G→A) nucleotide substitution and also displayed suppression. Together with our transgene rescue data, this confirmed the identity of the causal mutation in WY1211.

In the case of the dominant allele *fd132* (strain WY1210), we first engineered corresponding changes to the two candidates for strain WY1210 (T09B9.4, M399I; W07E11.1, P1181L). In the case of T09B9.4, we obtained >15 isolates but failed to observe suppression in any of these strains. Likewise, in the case of W07E11.1, we obtained more than five isolates, but these too failed to produce suppression. We also carried out dsRNA injections for both T09B9.4 and W07E11.1 into strains WY1145 and WY1210. If the *fd132* mutation is haploinsufficient or dominant negative, RNAi may be expected to cause suppression of WY1145 lethality. Alternatively, if the *fd132* mutation is gain-of-function or neomorphic, RNAi may be expected to revert suppression of WY1210. RNAi of both genes, however, failed to alter the phenotypes of either WY1145 or WY1210. Taken together, our data strongly indicate that the mutations in T09B9.4 and W07E11.1 do not correspond to the *fd132* causative mutation.

**Oligonucleotides**

For the following CRISPR targeting RNAs (crRNAs), target-specific sequences have a yellow background. For repair templates, altered nucleotides are in red, altered restriction endonucleases are underlined, and crRNA target sequences have a yellow background. For dsRNA template primers, target-specific sequences are underlined.

**C04A11.4 (WY1208; *fd130*)**

crRNA: 5'-CAGGAGUUCUGUUACGAAGGGUUUUAGAGCUAUGCU-3'
Alt-RTM CRISPR-Cas9 tracrRNA: IDT (CA#1072534) was used for all CRISPR injections.
Locus amplification: 5'-CCACCGAACAAACCCAATTG-3' and 5'-CGGAGCATTTAAGCCACCAT-3'
Locus sequencing: 5'-ATATTCGGTCATTGTGGCCC-3' and 5'-CTGGGTTTCACACTGCAACA-3'

**F56D12.6 (WY1209; *fd131*)**

crRNA: 5'-GCUCGAUCGGAAGCCGGCGCGUUUUAGAGCUAUGCU-3'
Locus amplification: 5'-GACTATCGCCTCAACTCAGA-3' and 5'-CTTCTCTATTTCCGCAGCTC-3'
Locus sequencing: 5'-GGAAGCGTGGATTCTGTGAT-3' and 5'-CAGCGAGCACCTTTTTACGA-3'
dsRNA primers: 5'-TAATACGACTCACTATAGGGAGAGGACGATTGGCGGAACAAAA-3' and
5'-TAATACGACTCACTATAGGGAGAGTGTCCAGGTTCGAATTTCC-3'

**B0302.1 (WY1217; *fd139*)**

crRNA: 5'-TGAGCCGATTCTCTCGTCTGGUUUUAGAGCUAUGCU-3'
Locus amplification: 5'-GCAACACGCCTATCACAGTT-3' and 5'-TCCGGAGAATCTGTTCTCTG-3'
Locus sequencing: 5'-TCAAATGTCGGACGAGGAGA-3' and 5'-CATTTAACTGAGGGTACCCG-3'
dsRNA primers: 5'-TAATACGACTCACTATAGGGAGACCATTCGAATATGCCCGCAA-3' and
5'-TAATACGACTCACTATAGGGAGATGAGTCACTGGTTTTGCCTG-3'

**F48E8.5 (WY1211; *fd133*)**

crRNA: 5'-GCAAAGAGTTTGAAGCGAATGUUUUAGAGCUAUGCU-3'
Locus amplification: 5'-GTTCTCAATCCGCGAGGCA-3' and 5'-TCGATTACAAGCCGAGTGCT-3'
Locus sequencing: 5'-GGAGCAGATTCTGAAGGAGA-3' and 5'-ACAGTTCTTGGCTTCCTCTG-3'
Repair template:
5'-CTTGTCGAGGATGACGTGCCGAATGTCAGATTCAACGCCGCAAAGAGTTTAAAGCGAA
TTGAAAAGAACTTGACCCCAAGGTGACAGGAAAATCTTTTTCACTATCCCAATTT-3'
(Creates DraI site)

**W07E11.1**
crRNA: 5'-TCAAACTTTGCACGGAGAACGUUUUAGAGCUAUGCU-3'
Locus amplification: 5'-CTCAGAAGCTGGAGTTGGAA-3' and 5'-TCTCTCCAGCAACTCCTCTA-3'
Locus sequencing: 5'-GTGCTCACGATGAACAACCT-3' and 5'-AATGGATCCGAACTCGAGCA-3'
Repair template:
GATGAGAAAATGTCACTTGAACACCTGCCCGGTTGGAGTCGCTACTCAAGATCTTGTTCTCCGTGCAAAG
TTTGATGGAAAGCCAGAACACGTTGTCAACTATATG
(Creates BglII site)
dsRNA primers:
5'-TAATACGACTCACTATAGGGAGAACTTAAAGTGCGCTAACCCG-3' and
5'-TAATACGACTCACTATAGGGAGACCACCAGAGAGACATTTTCC-3'

**T09B9.4**
crRNA: 5'-GCTTGTTGGAATGTGCAATCGUUUUAGAGCUAUGCU-3'
Locus amplification: 5'-ATCAGGAGGAATTGATGCGC-3' and 5'-CGGCTTCGATACGAATAGCT-3'
Repair template:
ATCGTCAGCAACTCAACGAAACTGCCCATTTGCACATTGAAAACAAAAAGCTAGTTGGAATATGCAATCA
AGAGAGACGTCGATCGCAAGCTACAGAGGCTGAAGTCAAAAAATgttagtga
(Ablates HindIII site)
dsRNA primers:
5'-TAATACGACTCACTATAGGGAGAACACCGCTGCCTGATAATCA-3' and
5'-TAATACGACTCACTATAGGGAGATGACTTCAGCCTCTGTAGCT-3'
