## Supplementary Materials for "Use of a Sibling Subtraction Method (SSM) for Identifying Causal Mutations in *C. elegans* by Whole-Genome Sequencing"

### Supplemental WGS and SSM Methods

***Strain Generation***

1) We recommend that mutant strains be backcrossed to remove most of the background mutations and to ensure that mutations of interest exhibit stable Mendelian properties. In our experience, backcrossing three to five times (with our starting strain) seems to be sufficient for SSM/WGS to work, and even fewer backcrosses may suffice. Backcrossing methods can also give important information as to whether mutations are dominant or recessive and whether they are linked to LGX (See Supplemental Genetics Methods).

2) As depicted in Figure 2 of the paper, heterozygous (e.g., *m/+*) hermaphrodites are allowed to self-fertilize, and progeny are cloned to individual plates to obtain the *m/m*, *m/+*, and *+/+* genotypes (1:2:1). Next, it is essential to definitively identify plates that are either *m/m* or *+/+*, which will be used to generate DNA for the Mutant and the Non-Mutant Sibling Comparator (NMSC) DNA pools, respectively. Do not use any isolate if there is doubt as to its genotype/phenotype. We were successful in obtaining data for SSM/WGS using as few as five isolates (*m/m* or *+/+*) for the Mutant and NMSC pools. In most cases, however, ~15 isolates were used. Depending on sequencing depth, greater numbers of isolates may work better.

3) Genomic DNA is next isolated from the *m/m* (Mutant pool) and *+/+ (*NMSC pool) isolates. For convenience, we generally pooled worms from five *m/m* or five *+/+* isolates per DNA sample. For example, in cases where we had obtained 15 isolates, we made three separate genomic DNA preps, each containing worms from five isolates. Equal molar proportions of these preps were then pooled (master pool) to provide DNA for sequencing.

***Preparation of Genomic DNA***

**Harvesting clean worm populations**

Solution:

M9

Procedure:

- Pick individual hermaphrodites (from the strain of interest) onto NGM plates spotted with OP50.
- Allow F1 and F2 progeny to develop until plates are crowded but have not been starved for more than a day. Always try to prepare 5–10 more plates than your desired number, as some plates may be contaminated, etc. In general, we prepare 20 plates for each Mutant or NMSC isolate.
- Not all plates will be ready at the same time. Wash the plates that are ready as follows:
    1. Wash each plate with 1 mL of M9 solution and transfer into individual 1.7-mL tubes.
    2. Spin them at 5 rpm for 1–2 min.
    3. Resuspend/rinse the worm pellet with M9.

4. Repeat Steps 2 and 3 three more times.
5. Add 1 mL of M9 and place 1.7-mL tubes on a shaker/mixer for 2 hr at room temperature to get rid of remaining bacteria in the gut.
6. Repeat steps 2 and 3 five additional times.
7. Spin the tubes at 5 rpm for 1–2 min and remove as much M9 as possible without disturbing the pellet.
8. Freeze the worms at –80°C overnight or until ready to use.

**Preparing genomic DNA**

Solutions:

- Gentra Puregene Tissue Kit (4 g)
  Cell lysis solution, Protein precipitation solution, DNA hydration solution, Puregene Proteinase K and RNase A Solution
- Isopropanol
- 70% Ethanol

Procedure:

1. Thaw the frozen 1.7-mL tubes from the five isolates and combine them into one 1.7-mL tube. (DO NOT add additional M9 to transfer the pellets).
2. Add 1 mL of Cell lysis solution and 5 µL of Proteinase K solution and incubate at 55°C for 3 hr or until worms are entirely lysed. (Invert worms periodically during incubation.) Allow the lysate to cool to room temperature.
3. Add 5 µL of RNase A solution and incubate at 37°C on a shaker/mixer for a minimum of 1 hr.
4. Cool on ice for 3 min.
5. Add 333 µL of Protein precipitation solution.
6. Cool on ice for 5 min.
7. Vortex vigorously at high speed for 20 seconds.
8. Cool on ice for 5 min.
9. Centrifuge at 21000 × *g* for 2 min, to precipitate the proteins.
10. Transfer equal volumes of the supernatant into two new 1.7-mL tubes.
11. Add 500 µL of isopropanol to each tube.
12. Mix by inverting 50 times.
13. Incubate at –20°C for 1 hr.
14. Centrifuge at 21000 × *g* for 5 min. A white pellet should be visible.
15. Remove the supernatant, add 500 µL of 70% ethanol to each tube and invert several times to wash the DNA pellet.
16. Centrifuge at 21000 × *g* for 5 min, remove the supernatant, and air-dry the pellet.
17. Add 30 µL of DNA hydration solution, pipette up and down, and transfer to a clean 1.7-mL tube.
18. Incubate at 65°C until the DNA is fully dissolved. Mix to distribute evenly.

19. Measure the DNA concentration using a NanoDrop or similar device.
20. Load 7–10 µL on an agarose gel. A major band of >10 kb should be observed. There should be no RNA bands.

**WGS, variant detection, and sibling subtraction method (SSM)**

Our sequencing effort was designed to provide 20× average coverage of the *C. elegans* genome (105 Mb) to provide adequate depth to detect causative single-nucleotide polymorphisms (SNPs) in our experimental datasets. 2×125 paired-end library construction (150-bp insert) and sequencing were performed using the Tru-Seq DNA PCR-free Library Preparation Kit, and all samples were sequenced in one lane on an Illumina HiSeq2000. We sequenced the NMSC **[genomic DNA]** for each mutant and also sequenced each of the five mutants described in this paper. All raw Illumina data are deposited in the NCBI Sequence Read Archive (http://trace.ncbi.nlm.nih.gov/Traces/sra) (accession number forthcoming). Quality assessment was carried out with FastQC (*http://www.bioinformatics.babraham.ac.uk/projects/fastqc/* ). Residual adapters and low-quality and short reads were removed via Trimmomatic v0.35 (Bolger *et al.* 2014).

Following quality control, we performed the remaining analyses with Galaxy (Afgan *et al.* 2016) following a modified version of the CloudMap protocol (Minevich *et al.* 2012). Protocols described below (*SSM Variant Detection* and *SSM Variant Subtraction*) can be accessed through the shared workflows on UseGalaxy.org. Individual tools used in Galaxy are noted in italics below (Li and Durbin 2009; Li *et al.* 2009; McKenna *et al.* 2010; Cingolani *et al.* 2012a; Cingolani *et al.* 2012b). Reads from each sample were aligned to *the C. elegans* WS220 (ce10)] genome with *Map with BWA for Illumina* [Galaxy version 1.2.3]. The parameters used for alignment were as follows: Library mate-paired = Paired-end, Maximum edit distance (aln -n) = 0, Fraction of missing alignments given 2% uniform base error rate (aln -n) = 0.04, Maximum number of gap opens (aln -o) = 1, Maximum number of gap extensions (aln -e) = -1, Disallow long deletion within [value] bp towards the 3'-end (aln -d) = 16, Disallow insertion/deletion within [value] bp towards the end (aln -i) = 5, Number of first subsequences to take as seed (aln -l) = -1, Maximum edit distance in the seed (aln -k) = 2, Mismatch penalty (aln -M) =3, Gap open penalty (aln -O) = 11, Gap extension penalty (aln -E) = 4, Proceed with suboptimal alignments if there are no more than INT equally best hits (aln -R) = Null, Disable iterative search (aln -N) = No, Maximum number of alignments to output in the XA tag for reads paired properly (samse/sampe -n) = 3, Maximum number of alignments to output in the XA tag for disconcordant read pairs (excluding singletons) (sampe -N) = 10, Maximum insert size for a read pair to be considered as being mapped properly (sampe -a) = 500, Maximum occurrences of a read for pairing (sampe -o) = 100000, Specify the read group for this file? (samse/sampe -r) = No, Suppress the header in the output SAM file = No, Job Resource Parameters = Use default job resource parameters.

Orphaned and misaligned pairs were removed with *Filter SAM* [Galaxy version 1.0.0], and then BAM file read groups were altered with *Add or Replace Groups* [Galaxy version 1.56.0] with the following parameters: Read group ID (ID tag) = 1, Read group sample name (SM tag) = rgSM, Read group library (LB tag) = rgLB, Read group platform (PL tag) = rgPU, Specify additional (optional) arguments = Use pre-set defaults, Output bam instead of sam = Yes.

Local realignment was carried out to reduce the number of false positives caused by indels in our experimental data as compared to the reference genome. Intervals for local realignment were identified with *Realigner Target Creator* [Galaxy version 0.0.4] using the default parameters in Galaxy: (1) Basic GATK options: How strict should we be in validating the pedigree information = STRICT, Interval set rule = UNION, Type of reads downsampling to employ at a given locus = NONE, Type of BAQ calculation to apply in the engine = OFF, BAQ gap open penalty (Phred Scaled) = 40, Use the original base quality scores from the OQ tag = No, Value to be used for all base quality scores, when some are missing = -1, How strict should we be with validation = STRICT, Interval merging rule = ALL, Disable experimental low-memory sharing functionality = No, Makes the GATK behave non deterministically, that is, the random numbers generated will be different in every run = No. (2) Basic Analysis options: Window size for calculating entropy or SNP clusters (windowSize) = 10, Fraction of base qualities needing to mismatch for a position to have high entropy (mismatchFraction) = 0.15, Minimum reads at a locus to enable using the entropy calculation (minReadsAtLocus) = 4, Maximum interval size = 500.

Local alignment was performed in sequence intervals identified above using *Indel Realigner* [Galaxy version 0.0.6], with the following parameters and Basic GATK and Basic Analysis options: LOD threshold above which the realigner will proceed to realign = 5.0, Use only known indels provided as RODs = No. **[1)** Basic GATK options: How strict should we be in validating the pedigree information = STRICT, Interval set rule = UNION, Type of reads downsampling to employ at a given locus = NONE, Type of BAQ calculation to apply in the engine = OFF, BAQ gap open penalty (Phred Scaled) = 40.0, Use the original base quality scores from the OQ tag = No, Value to be used for all base quality scores, when some are missing = -1, How strict should we be with validation = STRICT, Interval merging rule = ALL, Disable experimental low-memory sharing functionality = No, Makes the GATK behave non deterministically, that is, the random numbers generated will be different in every run = No. **(2)** Basic Analysis options: Percentage of mismatching base quality scores at a position to be considered having high entropy = 0.15, Simplify BAM = No, Consensus Determination Model = USE_READS, Maximum insert size of read pairs that we attempt to realign = 3000, Maximum positional move in basepairs that a read can be adjusted during realignment = 200, Max alternate consensuses to try = 30, Max reads (chosen randomly) used for finding the potential alternate consensus = 120, Max reads allowed at an interval for realignment = 20000, Don't output the original cigar or alignment start tags for each realigned read in the output bam = No.

Duplicate reads were then removed (*Mark Duplicate Reads* [Galaxy version 1.56.0]: Remove duplicates from output file = Yes, Assume reads are already ordered = Yes, Regular expression that can be used to parse read names in the incoming SAM file = [a-zA-Z0-9]+:[0-9]:([0-9]+):([0-9]+):([0-9]+).*, The maximum offset between two duplicate clusters in order to consider them optical duplicates = 100) and variants were identified with *Unified Genotyper* [Galaxy version 0.0.6] using the following parameters: Genotype likelihoods calculation model to employ = BOTH, The minimum phred-scaled confidence threshold at which variants not at 'trigger' track sites should be called = 30.0, The minimum phred-scaled confidence threshold at which variants not at 'trigger' track sites should be emitted (and filtered if less than the calling threshold) = 30.0. (1) Basic GATK options: How strict should we be in validating the pedigree information = STRICT, Interval set rule = UNION, Type of reads downsampling to employ at a given locus = NONE, Type of BAQ calculation to apply in the engine = OFF, BAQ gap open penalty (Phred Scaled) = 40.0, Use the original base quality scores from the OQ tag = No, Value to be used for all base quality scores, when some are missing = -1, How strict should we be with validation = STRICT, Interval merging rule = ALL, Disable experimental low-memory sharding functionality = No, Makes the GATK behave non deterministically, that is, the random numbers generated will be different in every run = No. (2) Basic Analysis options: Non-reference probability calculation model to employ = Exact, Heterozygosity value used to compute prior likelihoods for any locus = 0.001, The PCR error rate to be used for computing fragment-based likelihoods = 0.0001, How to determine the alternate allele to use for genotyping = DISCOVERY, Should we output confident genotypes (i.e., including ref calls) or just the variants? = EMIT_VARIANTS_ONLY, Compute the SLOD = No, Minimum base quality required to consider a base for calling = 17, Maximum fraction of reads with deletions spanning this locus for it to be callable = 0.05, Maximum number of alternate alleles to genotype = 5, Minimum number of consensus indels required to trigger genotyping run =5, Heterozygosity for indel calling = 0.000125, Indel gap continuation penalty = 10.0, Indel gap open penalty = 45.0, Indel haplotype size = 80, Vary gap penalties by context = No, Allow the discovery of multiple alleles (SNPs only) = No.

Variant call format (VCF) files for each sample were filtered using SnpSift [Galaxy version 1.0]; keeping only those variants with a read depth > 3 and that exhibit a 100% alternate allele frequency, ((GEN[0].AD[0] = 0) & (GEN[0].AD[1] > 3)). Subsequently, to filter only those NMSC variants that have a read depth > 3, the following filter expression was used: (DP>3). These filtered variant files were used for subtraction of NMSC variants from the mutant variants.

Subtraction was performed by implementing our SSM Variant Subtraction to remove any variants also found in the NMSC strain. For each sample, VCF files were subsetted using S*elect variants* [Galaxy version 0.0.3] with the following parameters and Basic GATK and Basic Analysis options: Don't include filtered loci in the analysis = No. **[(1)]** Basic GATK options: How strict should we be in validating the pedigree information = STRICT, Interval set rule = UNION, Type of reads downsampling to employ at a given locus = NONE, Type of BAQ calculation to apply in the engine = OFF, BAQ gap open penalty (Phred Scaled) = 40.0, Use the original base quality scores from the OQ tag = No, Value to be used for all base quality scores, when some are missing = -1, How strict should we be with validation = STRICT, Interval merging rule = ALL, Disable experimental low-memory sharing functionality = No, Makes the GATK behave non deterministically, that is, the random numbers generated will be different in every run = No. **[(2)]** Basic Analysis options : Don't update the AC, AF, or AN values in the INFO field after selecting = No, Output Mendelian violation sites only = No, Minimum genotype QUAL score for each trio member required to accept a site as a Mendelian violation = 0.0, Selects a fraction (a number between 0 and 1) of the total genotypes at random from the variant track and sets them to nocall = 0, Select only variants of a particular allelicity = ALL, Select a random subset of variants = Use all variants, Don't include loci found to be non-variant after the subsetting procedure = No.

Alternatively, the exact subtraction can be performed by the VCF-VCF intersect [Galaxy version 1.0.0_rc1.0] tool. Mutant variant file was selected as the second VCF dataset, and the NMSC variant file as the first VCF dataset. Then the following options were chosen to perform the subtraction: Union or intersection = Intersect, Invert selection? = Yes, compare records up to this many bp away (window size) = 30, output whole loci when one alternate allele matches = Yes, Advanced controls = Don’t use advanced options.

Finally, the subtracted variants were annotated using *SnpEff* [Galaxy version 1.0] with the following options: Input format = VCF, Output format = Tabular, Genome = *Caenorhabditis elegans*: WS220.64, Upstream/Downstream length = 10000 bases, Filter homozygous/heterozygous changes = No Filter, Filter homozygous/heterozygous changes = None, Chromosomal position = Use default (based on input type).

Results from the above analysis can be downloaded from Galaxy and further subsetted according to individual need, such as to include only those variants within coding regions or that occur on a particular chromosome. In addition, we recommend confirming candidate variants intended for experimental validation further by examining the individual NMSC and mutant alignments within a genome browser such as IGV (Robinson *et al.* 2011; Thorvaldsdóttir *et al.* 2013).

To relax the conditions for 100% alternative allele frequency in the SSM Variant Subtraction we have developed an alternative workflow, *SSM Variant Detection (Alternative)*. After removing the duplicate reads with *Mark Duplicate Reads* as described above, a pileup file was generated with MPileup [Galaxy version 2.1.3] (Koboldt, e*t al.*, 2012) with the following settings: Genotype Likelihood Computation = Do not perform genotype likelihood computation (output pileup), Output base positions on reads = No, Output mapping quality = No, Set advanced options = Basic, Set filter by flags = Do not filter, Select regions to call = Do not limit, Select read groups to exclude = Do not exclude, Disable read-pair overlap detection = No, Do not skip anomalous read pairs in variant calling = No, Disable probabilistic realignment for the computation of BAQ = No, Coefficient for downgrading mapping quality for reads containing excessive mismatches = 0, Max reads per BAM = 250, Redo BAQ computation = No, Minimum mapping quality for an alignment to be used = 0, Minimum base quality for a base to be considered = 13, Only generate pileup in region = NULL. The MPileUp outputs were subsetted with *Varscan* [Galaxy version 0.1]. *VarScan* allows for filtering variants across a range of allele frequency thresholds with SNPs and INDELs called separately. The SNP and INDEL VCF files obtained for each strain were combined using *VCFCombine* [Galaxy Version 1.0.0_rc1.0]. Finally, *SnpEff* was used to annotate variants with the parameters described above.

**Manual filtering of SSM/WGS candidates**

We find that it is essential to manually examine causal mutation candidates, as some false positives are likely to be produced by the bioinformatics pipeline. In our study, manual filtering decreased the total number of combined coding region candidates for the five suppressor strains from 32 to 11 (Table 1 and Supplemental File 1). In many cases, the false positives were variants that differed from the published N2 sequence but were present in 100% of Mutant and NMSC reads and thus represented background mutations in the starting strain. In several cases, variants were annotated as affecting coding regions but were in pseudogenes according to the current version of WormBase. We note that we did not detect any evidence for false negatives in our SSM/WGS dataset, although we cannot rule out this possibility. Our ability to identify all four of the recessive suppressor alleles strongly suggests that the frequency of false negatives is low. Moreover, in the case of the dominant allele *fd132*, current evidence points to this mutation affecting a non-coding region. Along these lines, a manual examination of reads within the 1,704,753 bp region on LGX (8889059–10593811) implicated to contain *fd132* did not produce any evidence of false negatives based on our analysis.

**EMS density mapping**

EMS density mapping was carried out as described (Zuryn *et al.* 2010). This method requires that background variants present in the starting strain be removed prior to EMS variant identification. To subtract background variants present in the starting strain, WY1208 was used to identify common variants also present in WY1209, WY1210, WY1211, and WY1217. For strain WY1208, common variants present in WY1209 were subtracted.
